# Bumblebees actively compensate for the adverse effects of sidewind during visually-guided landings

**DOI:** 10.1101/2022.12.16.520721

**Authors:** Pulkit Goyal, Johan L. van Leeuwen, Florian T. Muijres

## Abstract

Flying animals often encounter winds during visually guided landings. However, how winds affect their flight control strategy during landing is unknown. Here, we investigated how sidewind affects the landing strategy, sensorimotor control, and landing performance of foraging bumblebees (*Bombus terrestris*). For this, we trained a hive of bumblebees to forage in a wind tunnel, and used high-speed stereoscopic videography to record 19,421 landing flight maneuvers in six sidewind speeds (0 to 3.4 m s^−1^), which correspond to winds encountered in nature. Bumblebees landed less often in higher windspeeds, but the landing duration from free flight was not increased by wind. We then tested how bumblebees adjusted their landing control to compensate for the adverse effects of sidewind on landing. This showed that the landing strategy in sidewind was similar to that in still air, but with important adaptations. In the highest windspeeds, more hover phases occurred than during landings in still air. The rising hover frequency did not increase landing duration because bumblebees flew faster in between hover phases. Hence, they negated the adverse effects of increased hovering in high windspeeds. Using control theory, we revealed how bumblebees integrate information from the wind-mediated mechanosensory modality with their vision-based sensorimotor control loop. The proposed multi-sensory flight control system may be commonly used by insects landing in windy conditions and it may inspire the development of landing control strategies onboard man-made flying systems.

**Summary statement:** Bumblebees foraging in strong sidewinds can still land precisely on artificial flowers, allowing them to be efficient and robust pollinators in these adverse environmental conditions.

## Introduction

Wind is an important characteristic of the natural world that affects both ecological interactions and the biomechanics of flying insects. Wind affects their migration and dispersal (Hu et al., 2016; Mikkola, 1986), their interaction with plants and flowers (Alcorn et al., 2012), and floral visitation rates (Crall et al., 2020; Hennessy et al., 2021). Wind imposes maneuverability challenges (Burnett et al., 2020; Chang et al., 2016; Jakobi et al., 2018; Matthews and Sponberg, 2018; Mountcastle et al., 2015; Ortega-Jimenez et al., 2013; Ortega-Jimenez et al., 2014; Ravi et al., 2013; Ravi et al., 2016) that may increase flight-energetic costs (Combes and Dudley, 2009; Crall et al., 2017). Unravelling how flying insects cope with the effects of winds can help to understand their neuroethology, biomechanics, and ecology, as well as provide guiding principles for the development of wind mitigation strategies in man-made aerial vehicles.

Landing is an important behavior for all flying animals, in particular for animals such as bumblebees that rely on their landing ability to gather food essential for survival and reproduction (Goulson, 2010; Michener, 2007). Successful landing requires precise control of flight speed as an animal approaches the surface (Baird et al., 2013; Goyal et al., 2021; Srinivasan et al., 2000). While visiting flowers, foraging bumblebees land very frequently, with up to a thousand landings in an hour (Heinrich, 1979; Michener., 1974), and often in a wide range of wind conditions (Crall et al., 2017; Peat and Goulson, 2005; Riley et al., 1999).

In the absence of wind, many flying animals including bumblebees use visual feedback to control their flight speed as they advance towards the landing surface and achieve a soft touchdown (Baird et al., 2013; Balebail et al., 2019; Chang et al., 2016; Goyal et al., 2021; Lee et al., 1991; Lee et al., 1993; Liu et al., 2019; Reber et al., 2016; Srinivasan et al., 2000; Tichit et al., 2020; Van Breugel and Dickinson, 2012). Their motion relative to the landing surface generates optical expansion cues in which various features in the visual image appear to move radially outward from the point that is being approached (Edwards and Ibbotson, 2007; Gibson, 1955). Bumblebees can use this optical flow relative to the retinal image size of an object (Wagner, 1982), or the angular position of features in the image (Baird et al., 2013), to measure optical expansion rate, also known as relative rate of expansion. The instantaneous optical expansion rate *r* is equal to the ratio of approach velocity *V* of the bumblebee and its distance from the surface *y* (*r*=*V/y*; Figure 1A). Bumblebees use this optical expansion rate to control their landing (Chang et al., 2016; Goyal et al., 2021).

**Figure 1.**
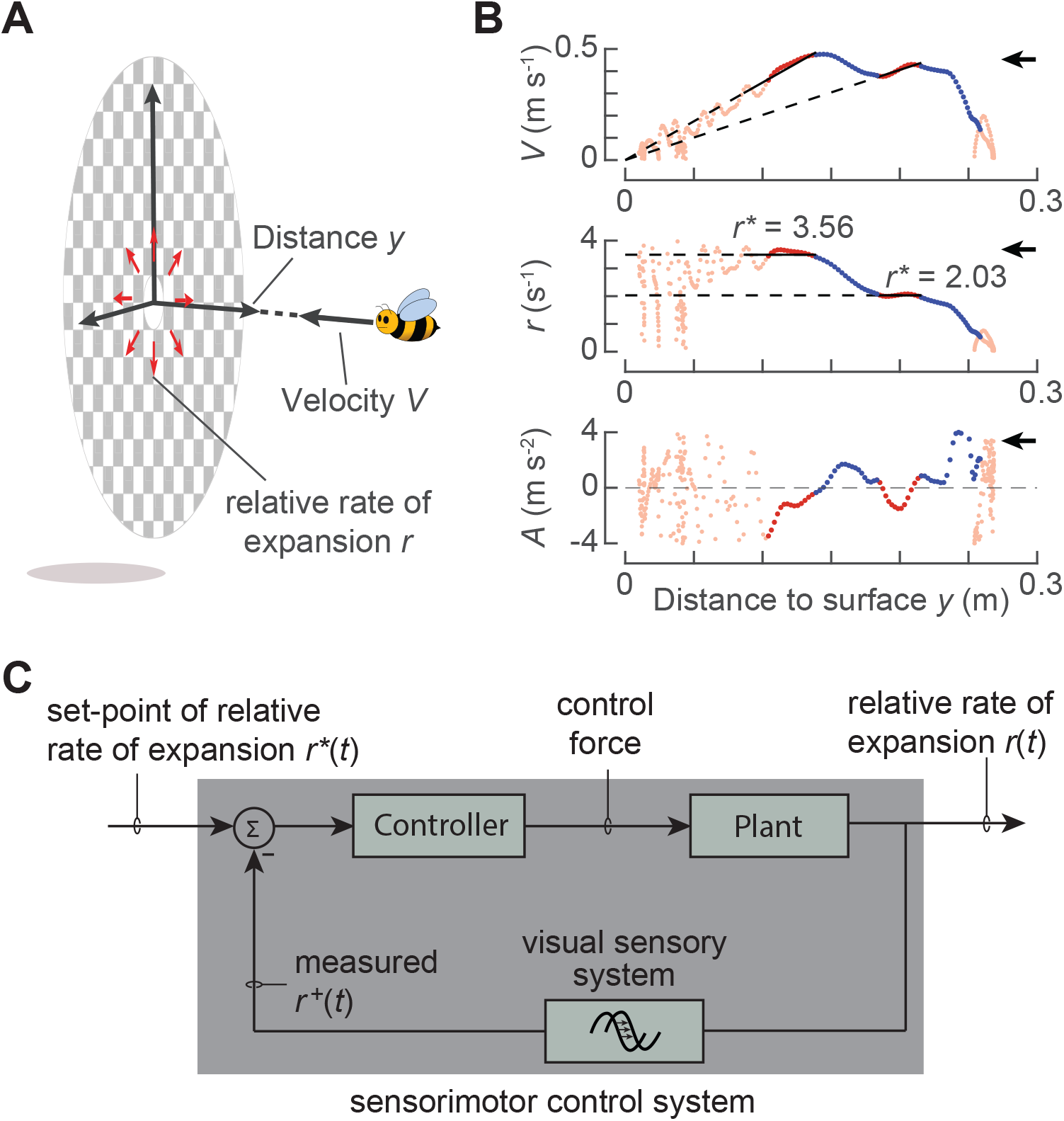
Visually-guided landing strategy and proposed sensorimotor control system of flying bumblebees. (A) Diagram of a bumblebee flying with perpendicular approach velocity *V* towards a landing platform). At a distance *y*, it experiences a relative rate of optical expansion *r* = *V/y*. (B) The bumblebee landing kinematics with plots of *V-y* (top), *r-y* (middle), and approach acceleration *A*=d*V*/d*t* versus *y* (bottom). The approach consists of an alternating series of entry segments (blue lines) and constant-*r* segments (red lines). Corresponding estimated set-points *r*\* are depicted by dashed black lines as slope and ordinate values in the *V-y* and *r-y* graphs, respectively. The black arrows indicate flight direction towards the landing surface. (C) Proposed closed-loop sensorimotor control system that landing bumblebees use to converge the optical expansion rate *r* to a set-point *r*\*. Using their visual system, bumblebees measure optical expansion rate as *r*^+^. Based on the difference between *r*^+^ and *r*\*, the animal produces a proportional aerodynamic control force, which accelerates the animal (“plant” in control terminology).

When doing so, bumblebees approach a landing surface in still air using a series of approach bouts (Figure 1B) (Goyal et al., 2021). During each bout, a bumblebee regulates the optical expansion rate *r*, and uses its sensorimotor control system to produce the motor output needed to reach a particular value of the optical expansion rate, also known as an optical expansion rate set-point *r*\* (Figure 1C) (Baird et al., 2013). As a result of these control actions, each flight bout consists out of two phases (Figure 1B) (Goyal et al., 2022). During the so-called constant-*r* phase, the bumblebee flies approximately at the optic expansion rate set-point, as regulated using steady-state control responses. Preceding the constant-*r* phase, bumblebees tend to use their transient flight responses to converge towards the constant-*r* set-points; these flight phases are defined as the entry segments. From one bout to the next, bumblebees tend to increase their set-point in a stepwise manner, leading to a new entry segment followed by a constant-*r* flight phase.

This stepwise upregulation of the optic expansion set-points causes landing bumblebees to intermittently increase their approach flight speed during the entry segments, and decrease it during the constant-*r* flight phase (Goyal et al., 2022). In addition to these accelerations and decelerations, bumblebees occasionally exhibit low velocity phases, also described as hover phases (de Vries et al., 2020; Goyal et al., 2021; Reber et al., 2016). These may result from an instability resulting from a flight controller that uses optical expansion rate as a control variable (Croon and De Croon, 2016).

During landing in wind, bumblebees experience different air speeds around their wings and body as compared to those in still air. As this airspeed influences the aerodynamic forces and torques that bumblebees produce with their flapping wings, it becomes mandatory for the bumblebees to adapt their sensorimotor control response for successful landings. This adaptation can be based on the active measurement of airspeed, possibly with their antennae (Jakobi et al., 2018; Taylor and Krapp, 2007), and must generate forces and torques that compensate for the effects of winds (Dickinson and Muijres, 2016; Dickinson et al., 2000).

In nature, winds are often characterized by a mean wind and the fluctuations around it (Garratt, 1994; Stull, 1988). While the effects of mean wind and the fluctuations on the locomotory performance have been studied in freely flying insects (Baird et al., 2021; Barron and Srinivasan, 2006; Crall et al., 2017; Engels et al., 2016; Fuller et al., 2014; Laurent et al., 2021; Ravi et al., 2015; Shepard et al., 2016), their effects on the landing behavior have received little attention. To our knowledge, only one study suggests that winds influence the landing dynamics of bumblebees (Chang et al., 2016), but it is unknown how bumblebees achieve flight control during these landing in winds. To address this knowledge gap, we here investigated the landing dynamics of bumblebees in the presence of various levels of steady sidewinds.

Specifically, we studied how a steady sidewind affects the vision-based modular guidance strategy and the sensorimotor control of landing bumblebees, and how bumblebees cope with these, potentially detrimental, effects. For this purpose, we exposed foraging bumblebees to six different steady horizontal winds ranging from 0 to 3.4 m s^−1^, directed parallel to the landing surface. These conditions correspond to the typical wind speeds that bumblebees experience in nature (Crall et al., 2017). Moreover, we applied sidewinds as bumblebees often encounter crosswinds during flight (Riley et al., 1999), and flying insects, including bumblebees, are most sensitive to the aerial disturbances along the lateral axis (Ravi et al., 2013; Ravi et al., 2016; Vance et al., 2013).

## Materials & Methods

### Experimental animals, setup and procedure

During our indoor experiments, lab temperature was maintained at 21±2°C. We used a hive with more than 50 female worker bumblebees (*Bombus terrestris*), provided by Koppert BV (Berkel en Rodenrijs, The Netherlands). The worker bees needed to perform forager flights to gather food from a 50% sugar solution, whereas dried pollen was provided in the hive.

Our experimental setup consisted of a wind tunnel (3.0×0.5×0.5 m; length×width×height), the bumblebee hive, a food source, and a real-time machine-vision-based videography system (Figure 2A). The hive and the food source were placed on two opposite sides of the middle section of the wind tunnel. Both were connected to the vertical walls of the wind tunnel (transparent polycarbonate 0.01 m thick sheets) using Plexiglass tubes (0.02 m diameter). These tubes were flush with the inside of the vertical tunnel walls. We attached circular landing platforms (0.18 m diameter) around each opening, with a visual pattern consisting of squares (1×1 mm) filled with random grayscale values (Figure 2B). The set-up was illuminated with a white broad-spectrum LED light panel to produce light intensities similar to overcast daylight conditions (1823 1x, Figure S1). For details see (Goyal et al., 2021).

**Figure 2.**
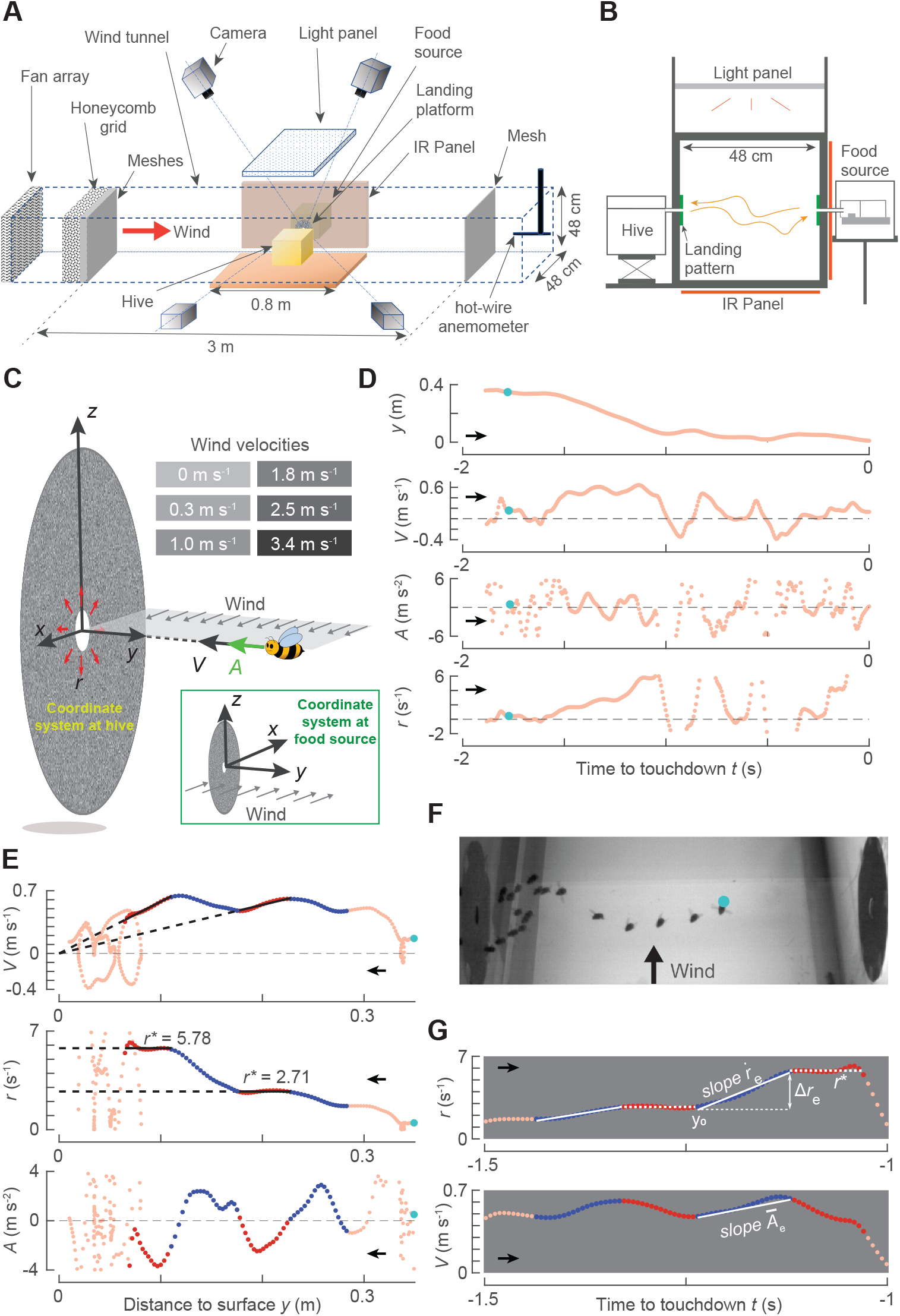
Experimental setup, flight kinematics during landing, and definitions of parameters. (A,B) The experimental setup consisted of a wind tunnel with two vertically placed circular landing surfaces connected to a hive and a food-source, respectively. Foraging bumblebees that flew between these landing surfaces were tracked in real-time using a four-camera videography system. Visible and IR LED light panel were used for background illumination and for videography illumination, respectively. (C) Landings on each platform were quantified in Cartesian coordinate systems, where the *x*-axis is parallel to the wind direction. (D–F) Landing maneuver of a free-flying bumblebee in a 3.4 m s^−1^ sidewind. In all panels, cyan dots denote the same instance, and black arrows show the approach flight direction. Parameters are approach distance *y*, velocity *V*, acceleration *A*, and relative rate of optical expansion *r*=*V/y*. (D) Temporal dynamics of (*y,V,A,r*), with *t*=0 s at touchdown. (E) Variation of (*V,r,A*) with *y*, where the constant-*r* and entry segments are in blue and red, respectively. (F) Photomontage from a top-view video at a time interval of ~0.1 s. (G) Definition of entry segment parameters: optical-expansion acceleration 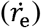, required stepchange in optical expansion rate (Δ*r_e_*), associated set-point (*r*\*), initial approach distance at entry segment start (*y*_0_), and average acceleration (*Ā*_e_).

To generate different steady wind conditions in the wind tunnel, we built two 6×6 grids of DC cooling fans (San Ace 80 9GA0812P7S001 or 9GV0812P4K03, Sanyo Denki Co., Japan). Both grids were powered with a 480 W power supply (Mean Well SP-480-12, Mean Well Co., Taiwan). The air flow generated by a fan-grid travelled through a honeycomb structure (Tubes core PC, diameter 6 mm, 100 mm thickness, Tubus Bauer GmbH, Germany) and a sequence of four meshes (FG1814F Fiberglass mosquito netting, 1.17×1.59 mm aperture, 68% transparency, Wire Waving Dinxperlo, The Netherlands) before it reached the wind tunnel test section. This was done to break down the fan-generated vortices and produce a low-turbulence uniform airflow.

We characterized the air flow in the wind tunnel using a hot-wire CTA anemometer (Dantec 55P16 wire probe and 54T42 MiniCTA, Dantec Dynamics, Denmark). We systematically measured the wind speed at various fan settings, days and locations within the tunnel. We first quantified the airflow the deviations from uniformity by measuring the airflow along the wind tunnel cross-section, in the middle of the wind tunnel, and at 0.20 m downstream and upstream locations. This showed that up to 4 cm from the walls the airflow speeds remain within 94% of the mean windspeed. Secondly, we measured the airflow variation over time for eleven consecutive days. Within this period, the windspeed varied maximally 2% from the mean windspeed. Thirdly, we used the hot-wire anemometer to quantify the turbulence intensity (standard deviation of the airflow speed divided by the mean) of the airflow in our set-up. This was less than 3% for all measured locations and for all tested wind speeds. These tests indicate that bumblebees experienced low-turbulent and close to uniform wind conditions, at distances of more than 0.04 m from the walls.

We investigated the landing dynamics of bumblebees in six steady wind speeds of 0, 0.3, 1.0, 1.8, 2.5 and 3.4 m s^−1^, which span the range of speeds that bumblebees commonly experience in nature (Crall et al., 2017). To generate these wind conditions, we controlled either of the two fan-grids using pulse-width modulation (the San Ace 80 9GA0812P7S001 fan-grid for 0.3 and 1.0 m s^−1^ winds, and the San Ace 80 9GV0812P4K03 fan-grid for 1.8, 2.5 and 3.4 m s^−1^ winds). For the zero-wind condition the fans were turned off.

Before starting the experiments, we trained the hive for four days to forage in still air. During training and experiments, bumblebees experienced a 10/14 hour day/night cycle. Each day, sunrise was simulated between 07:30h and 08:00h by gradually increasing the light intensity from zero to 1823 lx. Between 17:00h and 17:30h, we simulated sunset by gradually decreasing the light intensity back to zero. During these timeslots, bumblebees were exposed to zero wind condition. We divided the rest of the day (08:00h – 17:00h) into six 1.5-hour timeslots, and bumblebees were exposed to one wind condition in each timeslot following a pseudo-random schedule, spanned over eleven days (Table S1).

We used a customized machine-vision based videography system (Straw et al., 2011) to track in real-time (at 175 Hz) all three-dimensional flight movements in the wind tunnel test section (Figure 2A,B). The video system consisted of four high-speed cameras with a custom-built array of infrared LED panels for illumination (see (Goyal et al., 2021) for details). Based on the position coordinates of the tracked bumblebees, we reconstructed (and stored) the 3D flight trajectories of each bumblebee in a global Cartesian coordinate system, which was attached to the center of the specific landing surface, with the *y*-axis pointing normal the landing surface into the tunnel, the *z*-axis vertically upward and the *x*-axis in the downstream direction of the air flow (Figure 2C). Thus, different coordinate systems were defined at the hive side and at the food source. The coordinate system at the hive side is a right-handed system, whereas the system at the food-source is left-handed. This way, the wind always moved in the positive *x*-direction (Figure 2C).

### Estimation of state variables

We extracted all flight trajectories in which bumblebees landed on one of the landing platforms, using a previously-designed selection procedure (Goyal et al., 2021). These tracks were divided in landings on the landing platform at the food source or at the hive side. Additionally, we characterized the landing type of each track, being either a landing from free-flight or directly after taking off (from the ground of from the landing platform on the opposite side).

We filtered all tracks using a low-pass second-order two-directional Butterworth filter (cut-off frequency = 20 Hz, *filtfilt* in Matlab 2020a) and stored the filtered track as space-time arrays **X**=(*x,y,z,t*), with time *t* set to zero at the end of the landing maneuver (i.e., when a bumblebee was closest to the landing surface). We used a second-order central differentiation scheme to compute the corresponding velocity and acceleration arrays of the flying bumblebee. Both were defined in the landing-platform coordinate system as the ground velocity **U**^G^=(*u*^G^,*v*^G^,*w*^G^) and body acceleration vector **A**=(*a_x_,a_y_,a_z_*), respectively (Figure 2D).

In addition to the ground-velocity of the bumblebee **U**^G^, we also recorded at each time-step the wind-velocity **U**^W^=(*u*^w^,0,0) and the air-velocity of the bumblebee defined as **U**^A^=**U**^G^–**U**^W^=(*u*^A^,*v*^A^,*w*^A^)=(*u*^G^-*u*^w^,*v*^G^,*w*^G^). Here, *u*^w^ is the wind velocity in the landing-platform coordinate system. Vector **U**^A^ is the relative air velocity experienced by the bumblebee, ignoring the effect of the bumblebee itself on the airflow. Thus, **U**^A^ depends on both the wind velocity and the ground velocity of the bee, and therefore it changes with time. The magnitude of **U**^A^ is denoted as airspeed *U*^A^.

To describe the approach of bumblebees towards the landing surface, we computed the temporal dynamics of four state variables: approach distance from the surface *y*(*t*), approach velocity *V*(*t*)=−*v*^G^(*t*), approach acceleration *A*(*t*)=−*a_y_*(*t*), and the relative rate of optical expansion that a bumblebee experiences due to its motion normal to the landing surface *r*(*t*)=*V*(*t*)/*y*(*t*) (Figures 2D–2F). We used the velocity perpendicular to the surface for the computation of relative rate of expansion, as bumblebees landing in still air have been shown to progressively increase and decrease this component as they advance towards the landing surface (Goyal et al., 2021).

### Extraction and characterization of constant-r segments and entry segments

To determine if bumblebees in the presence of winds use a similar modular landing strategy as in still air (Goyal et al., 2021), we applied the same analysis approach as used in that study. The algorithm developed for this identifies track segments in which a bumblebee kept the relative rate of expansion nearly constant (called constant-*r* segments). The corresponding response is called the steady-state flight response. We characterize these constant-*r* segments with the average values of the state variables (*y*\*,*V*\*,*A*\*,*r*\*), where *r*\* is referred to as a set-point of relative rate of expansion (Figure 2E). It is an estimate of the *r*-value that a landing bumblebee aims to fly at using its sensorimotor control system (Goyal et al., 2021).

The set-point extraction algorithm used to identify the constant-*r* segments depends on a sensitivity factor *f* which restricts the variation allowed around the mean *r*\* for a segment to be identified as a constant-*r* segment. The sensitivity factor *f* characterizes the number of standard deviations σ around the mean *r*\* that are included in the set of constant-*r* segments. This algorithm uses generalized *t*-distributions. The sensitivity factor is multiplied by a scale parameter σ of these distributions to obtain the plausible intervals of variables that determine the constancy of *r* in a track segment (Goyal et al., 2021). Here, we present the results for sensitivity factor *f*=1, but our results remain similar for a wide range of sensitivity factors (0.25≤*f*≤2.5), albeit with variable numbers of identified constant-*r* segments.

To analyze the sensorimotor control response of bumblebees in different wind conditions, we used a second previously-developed algorithm (Goyal et al., 2022). This algorithm identifies the track segments that precede the constant-*r* segments, and that contain a monotonic variation (increase or decrease) of relative rate of expansion (Figure 2E). In still air, this monotonic variation of *r* is the transient response of the sensorimotor control system when converging to the optic expansion rate set-point (Goyal et al., 2022). We refer to these segments as entry segments, and the corresponding flight responses as transient responses. We characterize each entry segment with six state variables (Figure 2G): optical expansion-acceleration 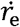, mean body acceleration *Ā*_e_, the required step-change in relative rate of expansion Δ*r*_e_, the associated set-point *r*\*, the initial approach distance at the start of the entry segment *y_0_*, and the mean airspeed during the entry segment 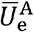.

Here, we use 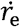 as a performance measure of the transient sensorimotor control response, as it dictates how fast a bumblebee reaches the expansion rate set-point *r*\*. For each entry segment, it is estimated from a linear regression: 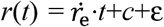, where *c* and ε denote intercept and residual, respectively. This linear regression captured well the motion during entry segments in all tested wind conditions (Figures 2G and 6A, Supplementary Material S3.1). Mean body acceleration *Ā*_e_ was computed as a ratio of change in approach velocity and travel-time during an entry segment. When *Ā*_e_>0, bumblebees accelerate towards the surface during the entry segment.

Our algorithms to extract the constant-*r* segments and entry segments do not capture all the set-points or entry phases that bumblebees exhibit during landing. For details about the limitations of each algorithm, see (Goyal et al., 2021) and (Goyal et al., 2022), respectively. We overcome these limitations by using thousands of landing maneuvers to describe the influence of winds on the landing dynamics of bumblebees.

### Characterization of the hovering phases exhibited by landing bumblebees

During landings, bumblebees may also rapidly break, causing them to greatly reduce their approach flight speed, and sometimes even hover or fly away from the landing surface (Figure 2E). These low-velocity flight phases are commonly described as hover phases (de Vries et al., 2020; Goyal et al., 2022; Reber et al., 2016), which we will do also.

To characterize how often bumblebees exhibited such hover phases during their landing in different sidewind speeds, we defined hover phases as sections during which the bumblebee reduced its approach flight speed *V* to below 0.05 m s^−1^. We then divided each landing trajectory into sections at four distances from the landing platform (0.05<*y*_1_<0.10 m, 0.10<*y*_2_<0.15 m, 0.15<*y*_3_<0.20 m, and 0.20<*y*_4_<0.25 m), and recorded the number of hover phases exhibited in each section. We did this for all landing maneuvers that started beyond section four (*y*=0.25 m). We applied a linear mixed-effects model to the resulting hover phase distributions, to test how the probability of exhibiting a hover phase *P*_hover_ varied among the four *y*-sections, with wind speed, and between the two landing types (landings after take-off and from free-flight).

### Quantification of the landing performance of bumblebees

To assess the overall landing performance of bumblebees at different sidewinds, we computed the travel time Δ*t* for each landing approach, which depicts how long bumblebees remain airborne during landing. The latter directly affects both the energetic cost and time budget of the landing maneuvers of foraging bumblebees. Therefore, minimizing Δ*t* can be a driving factor for maximizing landing performance. For all landing maneuvers that started beyond *y*=0.25 m, we computed Δ*t* as the time that a bumblebee takes when travelling from *y*=0.25 to *y*=0.05 m (distance 0.2 m). We then applied a generalized linear mixed-effects model to the travel time results, to test how Δ*t* varied with wind speed, and between landings directly after take-off and from free-flight.

### Statistical analyses

We used R 4.0.3 (The R Foundation, Austria) for statistical analyses. We developed linear mixed-effects models and a generalized linear mixed-effects model (using the R-functions *Imer* and *glmer*, respectively). Wherever relevant, we used the approach sequence, the landing side (whether a bumblebee landed on the side of the hive or food source), the day of the experiment, and the timeslot during the day as random intercepts. Probability values *p*<0.05 were considered statistically significant. For post-hoc comparisons, we used Bonferroni correction (using the *emmeans* package in R) to adjust the statistical significance values. The details of all developed models and associated results are found in Supplementary Material S3.2 and Tables S2-S8, respectively. We reported results and statistical estimates as mean values with either [95% confidence intervals], [standard error] or (standard deviation), depending on what is most appropriate.

## Results

We tracked the 3D flight kinematics of 19,421 landing approaches of bumblebees in the six sidewind speeds (*U*^W^=0 to 3.4 m s^−1^) (Table S1). Among these, 16,374 tracks represented landings from a free-flight, and 3,047 landings were performed directly after a take-off from the opposite landing platform or the ground. These landings resemble those when bumblebees move between flower patches and the hive, or when visiting multiple flowers within a single flower patch, respectively.

### Bumblebees land less often in high sidewind

We used two linear mixed models to test how the average landing frequency *N* varied with wind speed. The two models were separately applied to landings from a free-flight and after take-off (Supplementary Material S3.2; Figure 3; Table S2).

**Figure 3.**
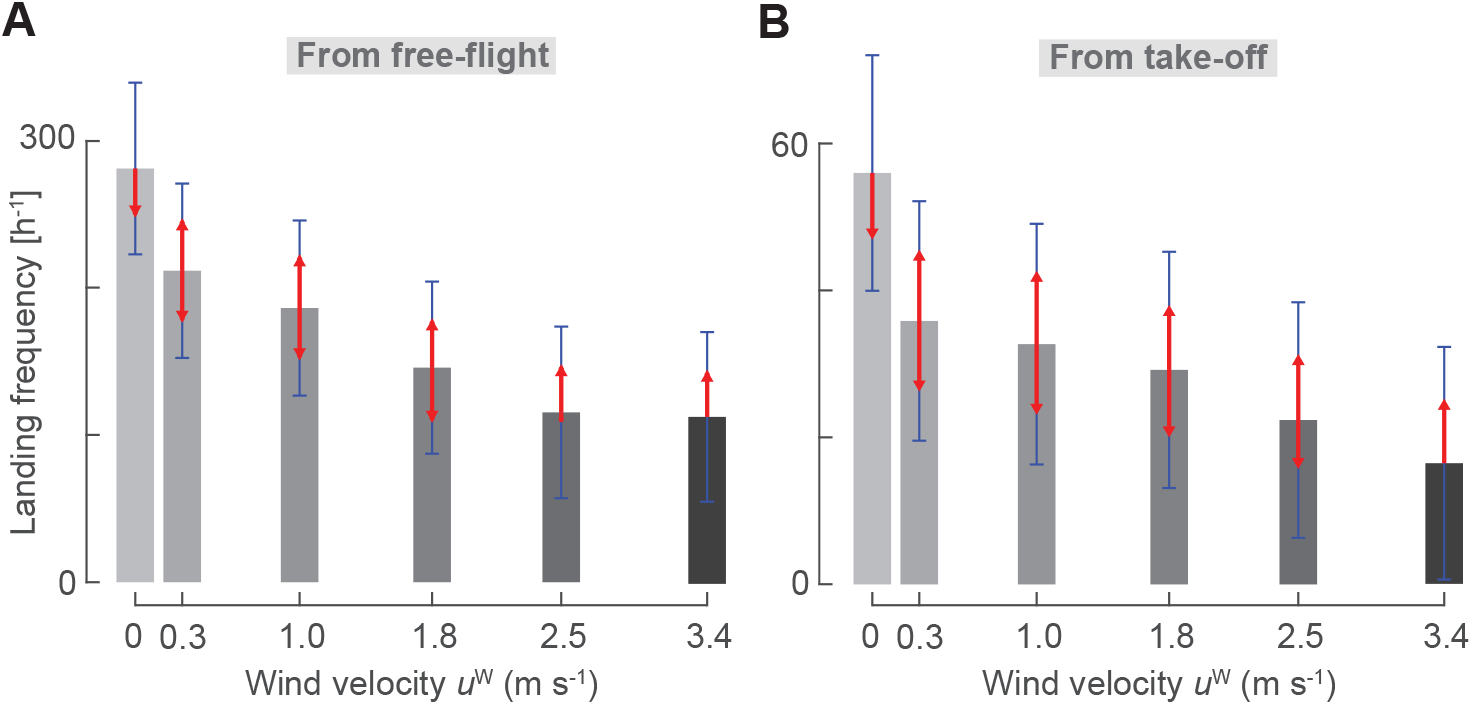
Bumblebees perform fewer landings at higher windspeeds. (A,B) Landing frequency, as landings per hour, in different sidewind velocities, for landings from free flight (A), or directly after take-off (B). Grey bars show the average landing frequency, blue bars show 95% confidence intervals, and non-overlapping red arrows indicate statistically significant differences between conditions (Table S2).

Landings from free-flight occurred 60% less in the highest winds (*U*^W^=3.4 m s^−1^) than in still air (*N*=112.2 [27.8] h^−1^ and *N*=280.8 [28.3] h^−1^, respectively; mean [standard error]; Figure 3A; Table S2). Landings after take-off occurred 70% less often in the highest wind condition (*U*^W^=3.4 m s^−1^) than in still air (*N*=16.5 [7.7] h^−1^ and *N*=56.0 [7.8] h^−1^, respectively; Figure 3B, Table S2). Thus, sidewinds reduce the landing frequency of foraging bumblebees.

### The average landing approach in different sidewind speeds

In all tested sidewinds, bumblebees flew on average approximately perpendicular to the landing surface (Figure 4A,B). Bumblebees experienced higher airspeeds *U*^A^ in higher sidewinds (Figure 4C,D). Thus, they had to generate higher compensatory sideways forces and torques during their landing approach. Most importantly, they needed to compensate for the additional drag force in the lateral direction due to winds. On average, they did this in all tested wind conditions, though with a slight lateral drift in the wind direction and a small height loss (Figure 4A,B).

**Figure 4.**
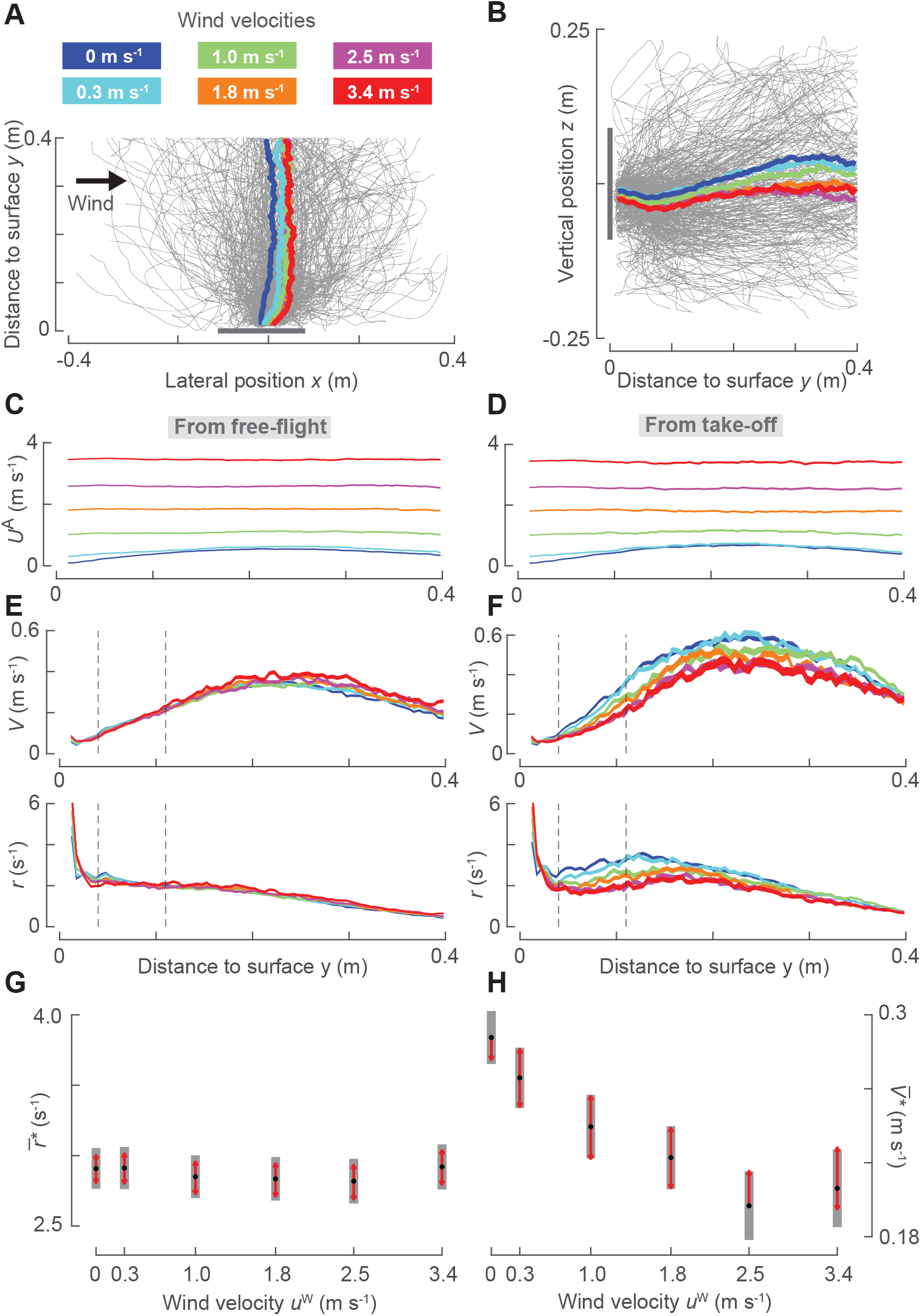
Effect of sidewind on the average approach kinematics of landing bumblebees. (A,B) Top and side views of landing maneuvers of bumblebees in all tested wind conditions. Grey lines show every 70^th^ recorded landing maneuver (*n*=281 tracks). Colored lines show mean landing maneuvers in different wind conditions (legend in A). (C-H) Differences in landing kinematics with wind speed (color-coded as in (A)); left and right panels show results for landings from free-flight and after take-off, respectively. The kinematics parameters are mean airspeed *U*^A^ (C,D), approach speed *V* and optic expansion rate *r* (E,F); line thickness indicates the standard error of the means. (G,H) Linear mixed model estimates of the average relative rate of expansion set-point 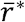 and resulting average approach velocity 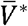 at *y*=0.075 m (Table S3). Black dots depict estimated means, grey bars show 95% confidence intervals, and non-overlapping red arrows indicate significant differences between conditions. Two vertical dashed grey lines in (E,F) show the distance range (0.04<*y*<0.11 m) for modelling 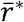.

On average, bumblebees first gradually increased and then decreased their approach velocity *V* as they approached the landing surface (Figure 4E,F). During their deceleration phase (0.04<*y*<0.11 m), the bumblebees flew on average at a nearly constant average set-point of optical expansion rate 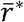 – a behavior described previously for honeybees (Baird et al., 2013; Srinivasan et al., 2000) and bumblebees (Chang et al., 2016; Goyal et al., 2021). However, individual flight paths can deviate substantially from this average behavior (Goyal et al., 2021).

We used a linear mixed model to find how this average set-point 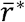 during the deceleration phase of the landing maneuver varied between wind conditions and landing types (from free-flight or take-off) (Supplementary Material S3.2). During landing from free-flight, bumblebees had similar average set-points in all wind conditions 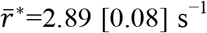 at 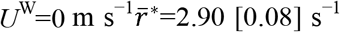 at *U*^W^=3.4 m s^−1^; Figure 4E,G; Table S3), and thus similar approach velocities throughout the deceleration phase.

However, when bumblebees landed shortly after a take-off, they decreased their average set-point with increasing wind speed (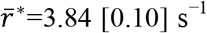 at *U*^W^=0 m s^−1^ and 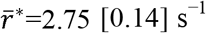 at *U*^W^=3.4 m s^−1^; Figure 4F,H; Table S3). Hence, only at low wind speeds (<1.7 m s^−1^) landings from take-off had higher set points than landings from free-flight; at high sidewinds the set points were similar (Figure 4G,H).

### At all windspeeds, landing bumblebees stepwise modulated their optical expansion set-point

The analysis of the average of landing maneuvers does not capture the detailed landing dynamics of individual bumblebees in still air (Goyal et al., 2021). Therefore, to understand how bumblebees land in sidewinds, we also analyzed the individual landing maneuvers using a previously-developed analysis approach (Figure 5A,B) (Goyal et al., 2021; Goyal et al., 2022).

**Figure 5.**
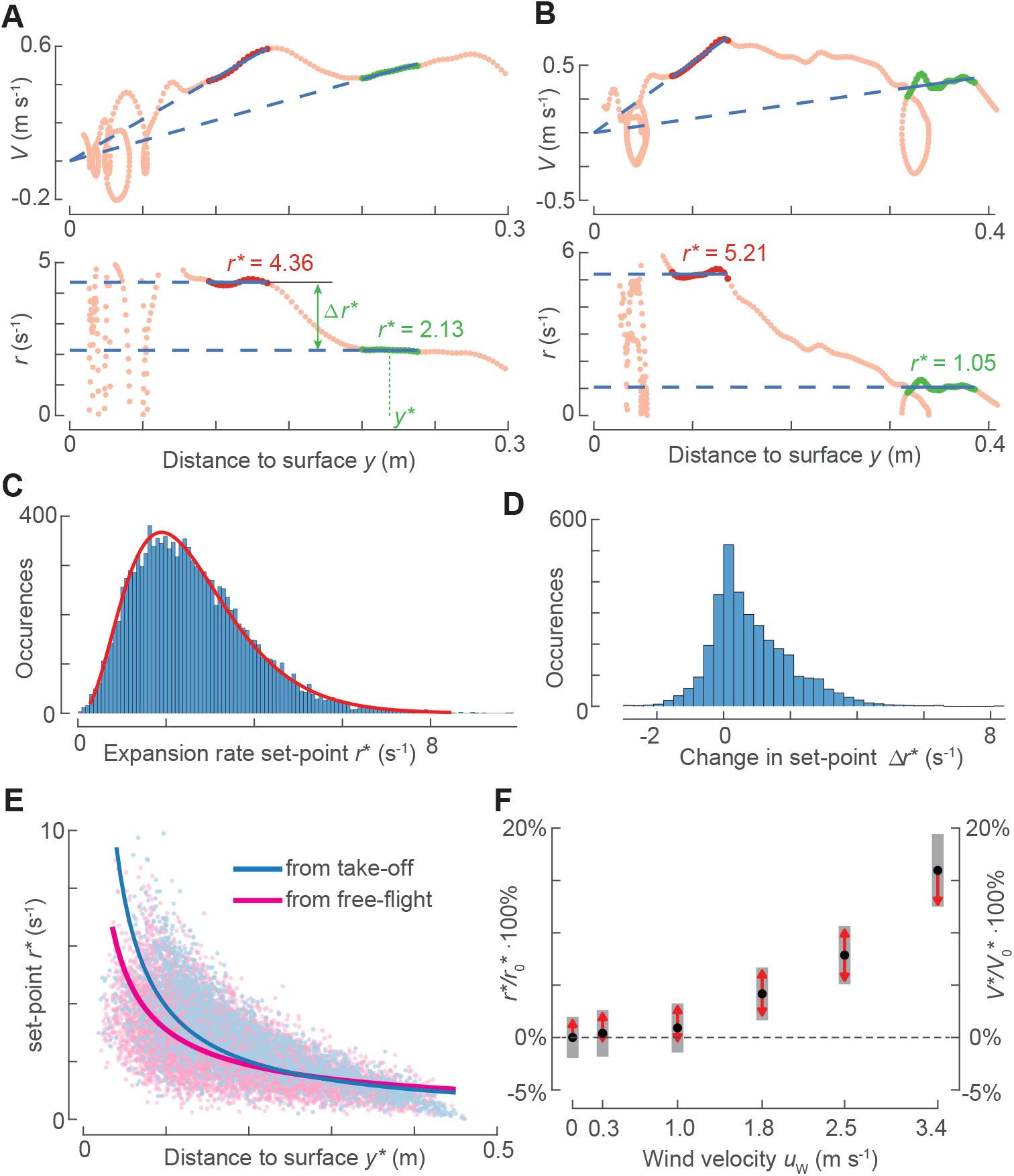
Bumblebees landing in a sidewind fly at higher optical expansion set-points than those landing without wind. (A,B) Approach velocity *V* and relative rate of expansion *r* versus distance *y* for bumblebees landing in a 3.4 m s^−1^ sidewind, after a take-off (A) and from free-flight (B). Two constant-*r* segments (green and red, respectively) and their optical expansion rate set-points *r*\* (blue) are highlighted. (C) Histogram of all identified set-points *r*\* (*n*=12,338), with gamma distribution fit in red. (D) Histogram of change Δ*r*\* between two consecutive constant-*r* segments in a landing maneuver, as defined in (A) (*n*=3,241). (E) Variation of *r*\* with corresponding distance *y*\* (defined in (A)), for landings after take-off (blue) and from free-flight (pink). Solid lines depict output of linear mixed models (Table S4). (F) Effect of wind on *r*\* and *V*\*, defined as percentage change relative to values at zero wind (*r*_0_* and *V*_0_*). Black dots depict estimated means, grey bars show 95% confidence intervals, and non-overlapping red arrows indicate significant differences.

Using a set-point extraction algorithm (Goyal et al., 2021), we identified 12,338 constant-*r* segments in 9,097 landing tracks (for sensitivity factor *f*=1) (Figure 5C). We approximated the observed distribution of identified set-points of relative rate of expansion *r*\* in these constant-*r* segments by a gamma distribution (median *r*\*=2.41 s^−1^, *a*=3.74 [3.65–3.83], *b*=0.69 [0.68–0.71], mean [95%confidence interval]) (Evans et al., 2000). This gamma distribution resembles the one found for bumblebees landing in still air (Goyal et al., 2021).

Out of the 9,097 landing tracks with constant-*r* segments, 2,632 had more than one constant-*r* segment (see Figure 5A,B for examples). In these tracks, bumblebees switched from one set-point to another 3,241 times, which occurred in all wind conditions (Figure 5D). In 76% of these 3,241 set-point transitions, bumblebees switched to a higher set-point resulting in a set-point increase of Δ*r*\*=1.24 (1.09) s^−1^ (mean (standard deviation)). For the remaining 24% of the transitions, bumblebees reduced their set-point with Δ*r*\* = −0.48 (0.48) s^−1^. These transition dynamics resemble those of bumblebees landing in quiescent air (Goyal et al., 2021). Thus, also in sidewinds, landing bumblebees switch more often to a higher set-point than to a lower set-point.

In the 9,097 landing tracks, we then tested how bumblebees adjusted their set-point of relative rate of expansion with distance to the surface. A linear relationship occurred between the logarithmic transformations of *r*\* and the corresponding distance to the surface *y*\* (Figures 5E and S2, Supplementary Material S3.2, Table S4). We used a linear mixed-effects model to find an estimate of the slope *m* of this linear fit. The model predicted that bumblebees, on average, increased their set-point with decreasing distance to the surface at a rate *m*=–0.727 [0.008] and *m*=–0.960 [0.017] while landing from free-flight and take-off, respectively. Surprisingly, these dynamics were independent of wind speed (Tables S4).

### In higher windspeeds, bumblebees approach the landing surface faster and at higher optic expansion set-points

Although *m* did not vary significantly with wind speed, the baseline optic expansion set-points at which bumblebees land in a sidewind were higher than for landings in still air (Figure 5F). This increase in set-point with wind occurred at all distances to the surface, for landing from both free-flight and take-off. This effect was well captured by a linear regression (Figure 5F, Table S4). At sidewinds of 2.5 m s^−1^ and 3.4 m s^−1^, the regression fits were *r*_2.5_* =1.08·*r*_0_* and *r*_3.4_* =1.16·*r*_0_* (subscript denotes the wind speed *U*^W^), and thus at these speeds bumblebees flew on average at an 8% and 16% higher set-point than in still air, respectively. Hence, bumblebees exhibited higher set-points in faster winds, and thereby flew faster towards the surface in higher windspeeds.

### Bumblebees exhibit faster sensorimotor control responses in higher windspeeds

The stepwise modulation of the set-point of optical expansion rate results in entry track segments that precede these constant-*r* segments, and contain the transient response of the sensorimotor control system (Figure 2E,G). We extract and analyze these entry segments using a previously-developed approach (Goyal et al., 2022) (see Materials & Methods for details). The entry-segment extraction algorithm linked 4,374 constant-*r* segments with a respective entry segment (see Figures 1B, 2E,G and 6A,B for examples). We modelled the sensorimotor control response of bumblebees during these segments as a motion at a constant optical-expansion-acceleration 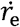 (Figure 6A) (see Materials & Methods); 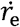 defines how fast a bumblebee reaches its set-point.

**Figure 6.**
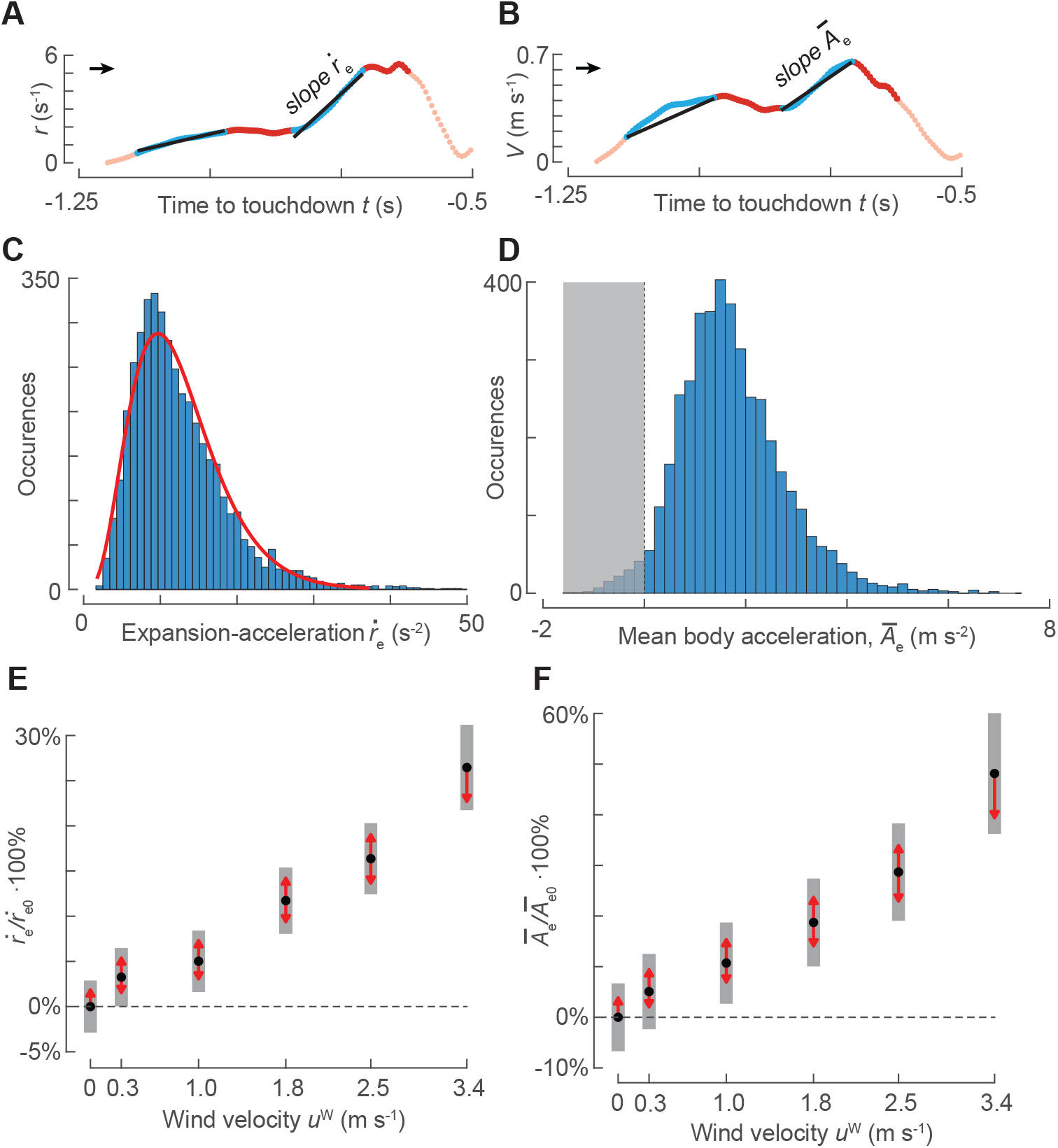
In sidewind, bumblebees accelerate faster towards the landing platform during the entry segments than in landings without wind. (A,C,E) Effect of sidewind speed on expansion-acceleration 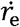 during the entry segments. (B,D,F) Effect of sidewind speed on accelerations towards the landing platform *Ā*_e_ during entry segments. (A,B) Temporal dynamics of optic expansion rate *r* and approach velocity *V*, including acceleration parameter definitions. (C,D) Histograms of occurrences of 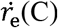(C), and *Ā*_e_ (D), for all identified entry segments with increasing optical expansion rate (*n*=4,221). (C) Red line shows the fitted gamma distribution. (E,F) Effect of wind on 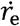 (E) and *Ā*_e_ (F), defined as percentage change relative to values at zero wind (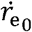 and *Ā*_e0_). Black dots depict estimated means, grey bars show 95% confidence intervals, and non-overlapping red arrows indicate significant differences (Tables S5,S6).

Among the 4,374 identified entry segments, bumblebees increased and decreased their optical expansion rate 4,221 and 153 times, respectively. Because in 97% of all identified entry segments, bumblebees increased their optical expansion rate, we focus only on analyzing the sensorimotor control response in these. In the 4,221 entry segments, the observed distribution of expansion-acceleration 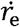 in the presence of all winds can be approximated by a gamma distribution (median 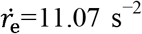, *a*=4.55 [4.4–4.7], *b*=2.7 [2.6–2.9]) (Figure 6C) (Evans et al., 2000). This distribution resembles the corresponding one identified for bumblebees in still air (Goyal et al., 2022).

We further used a linear mixed model to test how the observed optical-expansionacceleration 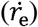 during an entry segment varied with wind speed. In addition to wind, this model also had other co-variates associated with an entry segment: *y*_0_, Δ*r*_e_, *r*\* and landing type (landing from free-flight or take-off) (Figure 2G, see Materials & Methods for their definition). The variations in expansion-acceleration 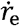 with these co-variates (Table S5) was found to be similar to those in still air (Goyal et al., 2022). Therefore, we focus on the effect of sidewind on the optical-expansion-acceleration 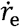 only.

Our model predicts that bumblebees increased the optical-expansion-acceleration with wind velocity, and thus landing bumblebees reached their set-points faster at higher windspeeds (Figure 6E). For example, in a 2.5 m s^−1^ and 3.4 m s^−1^ sidewind, bumblebees reached their set-point 16% and 27% faster than in still air, respectively (Figure 6E, Table S5). This wind effect was observed independently of all covariates (*y*_0_, Δ*r*_e_, *r*\* and landing type). Hence, winds augmented the transient response of the sensorimotor control system of landing bumblebees, allowing them to reach their set-point faster in higher sidewinds.

### Bumblebees accelerate faster towards the landing surface in higher windspeeds

Previously, we showed that in still air bumblebees mostly use the transient response of their sensorimotor control system to accelerate towards the landing surface (Goyal et al., 2021; Goyal et al., 2022). To understand how bumblebees use this transient response in the presence of winds, we computed the mean body acceleration *Ā*_e_ in all 4,374 identified entry segments (see example in Figures 2G and 6B).

In 153 of these 4,374 entry segments, bumblebees reduced their approach speed (*Ā*_e_<0) to decrease their optical expansion rate 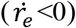. In the remaining 4,221 entry segments, bumblebees accelerated on average towards the landing surface (*Ā*_e_>0). In 4,102 entry segments, this led to an increase in the optical expansion rate 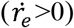, and in 119 entry segments it weakly decreased 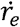 (Figure 6D). Hence, in 94% of all identified entry segments, bumblebees landing in a sidewind used the transient response of their sensorimotor control system to accelerate towards the landing surface. Moreover, the body accelerations produced by the bumblebees during these entry segments correspond mostly to an increase in relative rate of expansion.

Using a linear mixed model, we further tested how the positive average body acceleration *Ā*_e_ varied with sidewind velocity; this model includes the covariates (*y*_0_, Δ*r*_e_, *r*\* and landing type). The effects of these covariates on acceleration *Ā*_e_ in sidewinds were similar to those in still air (Table S6) (Goyal et al., 2022), and thus we will focus on the effect of sidewind on *Ā*_e_.

Our model predicts that bumblebees accelerated faster towards the landing surface in higher sidewind speeds (Figure 6F). For example, in a 2.5 m s^−1^ and 3.4 m s^−1^ sidewind, bumblebees accelerated on average 29% and 48% faster towards the landing platform than in still air, respectively. This behavior was independent of all covariates (*y*_0_, Δ*r*_e_, *r*\* and landing type, Table S6).

### Bumblebees hover more often with higher windspeeds and decreasing distance to the landing surface

Bumblebees approaching a landing surface in still air occasionally exhibit moments of near-zero or negative approach velocities (Figure 7A) (de Vries et al., 2020; Goyal et al., 2021; Reber et al., 2016). We will call these low velocity flight phases hover phases, as done in literature. These phases are potentially unfavorable as they tend to increase landing duration, which is energetically costly (Reinhold, 1999), in particular in the presence of winds (Shepard et al., 2016). Moreover, for foraging bees, an increase in landing time can reduce their floral visitation rate, and hence their energy gain (Hansen et al., 2002; Roubik, 1978).

**Figure 7.**
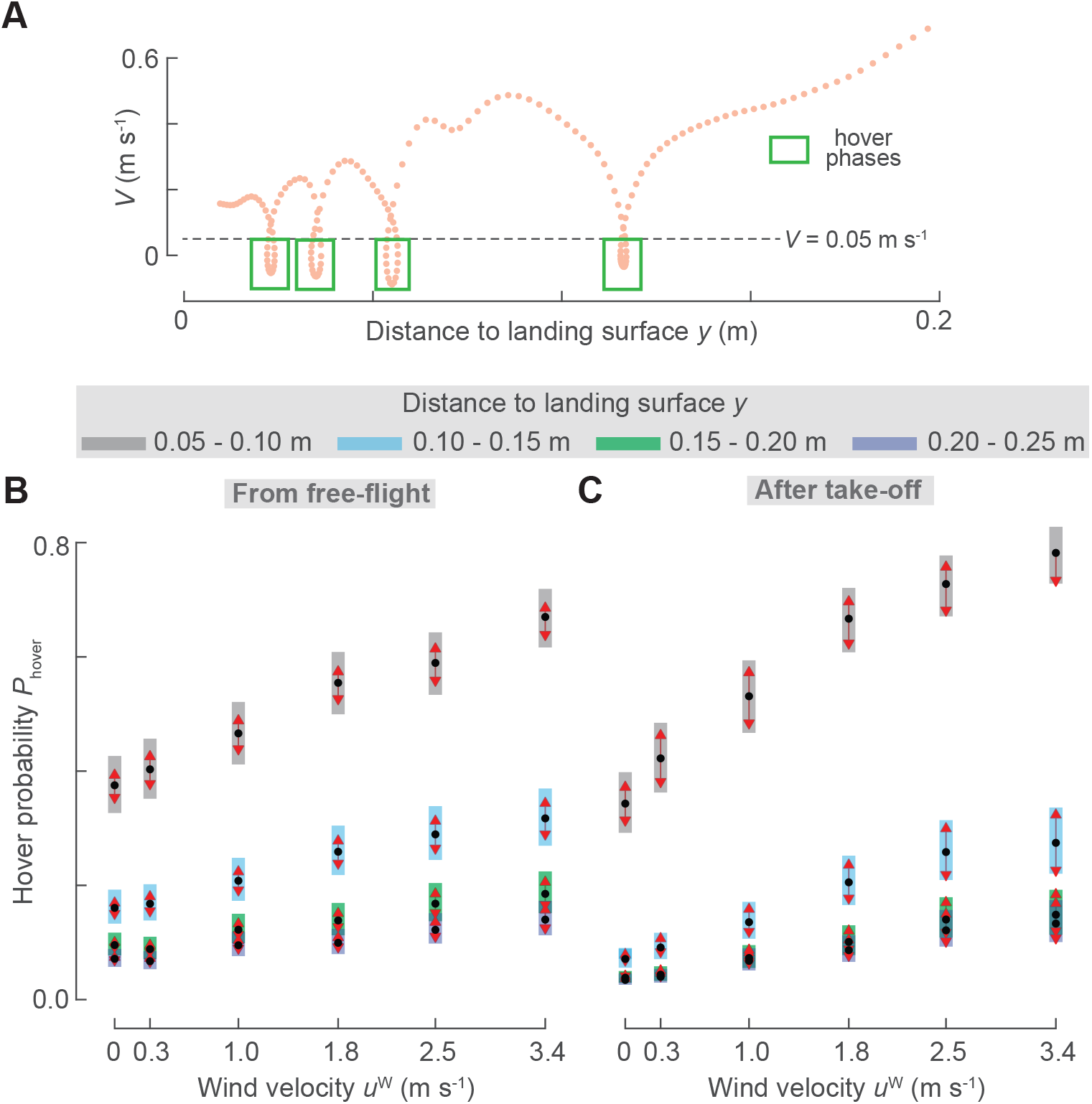
Bumblebees generate more hover phases when flying in higher sidewinds. (A) Approach velocity *V* versus distance *y*, with each hover phase (*V*<0.05 m s^−1^) highlighted with a green box. (B,C) Estimated probabilities that a bumblebee transitions to a hover phase at different wind speeds (abscissa), and different distances *y* (color-coded), for landings from free-flight (B) or after take-off (C). Black dots depict estimated means, grey bars show 95% confidence intervals. Non-overlapping red arrows indicate significant differences (Tables S5-S7).

We identified hover phases during a landing maneuver as track segments during which the approach speed dropped below 0.05 m s^−1^ (*V*<0.05; Figure 7A). We then used a generalized linear mixed model to test how the probability of exhibiting a hover phase *P*_hover_ varies with wind speed, landing type (landing from free-flight or take-off) and distance to the surface *y*. We divided the distance to the surface into four regions (0.05<*y*_1_<0.10 m, 0.10<*y*_2_<0.15 m, 0.15<*y*_3_<0.20 m and 0.20<*y*_4_<0.25 m) (Supplementary Material S3.2).

Bumblebees exhibited more hover phases in faster winds, for both landing types (landing from free-flight and take-off) and at all distances from the surface (Figure 7B,C; Table S7). Moreover, all explanatory variables had statistically significant two-way interactions (see below).

#### The effect of distance and landing type on hover probability

For landings from both free-flight and take-off, hovering probability increased with decreasing distance to the platform, but this distance effect on hovering probability was 65% larger in landing from take-off than in those from free-flight (Figures 7B,C). For landings initiated directly after take-off, hover probability was more than eight times higher closest to the platform (0.05<*y*_1_<0.10 m) than at the furthest analyzed distance (*P*_hover_=0.59 [0.03] for 0.05<*y*_1_<0.10 m and *P*_hover_=0.07 [0.01] for 0.20<*y*_4_<0.25 m, results are averaged over windspeeds). For landing from free flight, the equivalent hover probabilities differed only by a factor five (*P*_hover_=0.51 [0.03] for 0.05<*y*_1_<0.10 m, and *P*_hover_=0.10 [0.01] for 0.20<*y*_4_<0.25 m).

#### The effect of distance and windspeed on hover probability

At all distances from the landing platform, hovering probability increases with sidewind speed (Figure 7B,C; Table S7). This effect of sidewind on hovering probability depends significantly on the distance from the landing platform. For example, at the furthest tested distance (0.20<*y*_4_<0.25 m), hover probability in the fastest tested windspeed was 2.8 times that in still air (*P*_hover_=0.14 [0.02] at *U*^W^=3.4 m s^−1^, and *P*_hover_=0.05 [0.01] at *U*^W^=0 m s^−1^, results are averaged over landing types). At the closest tested distance (0.05<*y*_1_<0.10 m), hover probability increased only a factor 2 from no wind to highest windspeed (*P*_hover_=0.73 [0.02] at *U*^W^=3.4 m s^−1^, and *P*_hover_=0.36 [0.02] at *U*^W^=0 m s^−1^).

#### The effect of landing type and windspeed on hover _ probability

Finally, hover probability consistently increases with sidewind speed, for both landings from free-flight and from take-off (Figure 7B,C; Table S7). But wind affected hovering probability more for landings directly after take-off than for landings from free-flight. After take-off, the hover probability in the fastest tested windspeed was 3.8 times the probability in still air (*P*_hover_=0.30 [0.03] at *U*^W^=3.4 m s^−1^, and *P*_hover_=0.08 [0.01] at *U*^W^=0 m s^−1^, results are averaged over all landing distances). In contrast, after free-flight, the equivalent hover probability ratio was only 2 (*P*_hover_=0.30 [0.02] at *U*^W^=3.4 m s^−1^, and *P*_hover_=0.15 [0.01] at *U*^W^=0 m s^−1^). Hence, for landings after take-off the effect of wind on hovering probability is on average 88%larger than for landings from free-flight.

These combined results show that landing bumblebees generated more hover phases when they encountered higher sidewind speeds, particularly so directly after take-off (Figure 7).

### For landings from free-flight, travel time does not vary with sidewind speed

The increasing hover frequency at higher wind velocities might affect the travel time Δ*t* during the landing approaches of bumblebees. We tested this by calculating for each landing approach the time it took the bumblebee to travel from a distance of 0.25 m from the landing platform to a 0.05 m distance. These are the bounds of the four distance bins *y*_1_ to *y*_4_. We then used a linear mixed model to determine how Δ*t* varied with sidewind speed, and between landing from free-flight and take-off (Figure 8; Table S8).

**Figure 8.**
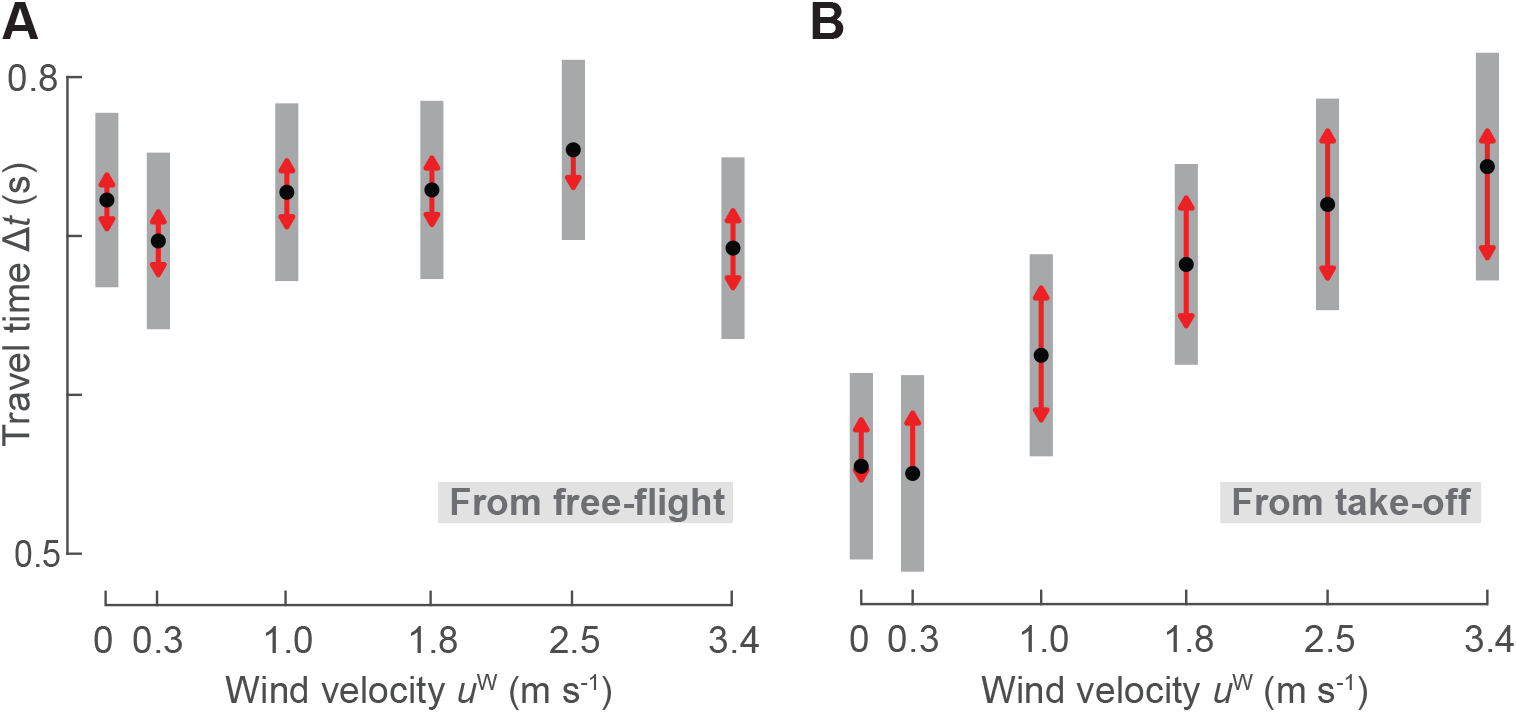
Travel time of bumblebees landing from free-flight does not vary with windspeed, whereas for landings after take-off travel time increase with sidewind speed. (A,B) Effect of sidewind speed on travel time Δ*t* defined as the time needed to travel from *y*=0.25 m to *y*=0.05 m, for landings from free-flight (A) or after take-off (B). Black dots depict estimated means, grey bars show 95% confidence intervals. Non-overlapping red arrows indicate significant differences (Table S8).

Surprisingly, for landings from free-flight travel time did not differ significantly between windspeeds, including landings in the highest wind speed and in still air (Δ*t*=0.69 [0.03] s at *U*^W^=3.4 m s^−1^, and Δ*t*=0.72 [0.03] s at *U*^W^=0 m s^−1^). However, the travel time after a take-off was 35% higher in the strongest sidewind as compared to those in still air (Δ*t*=0.74 [0.04] s at *U*^W^=3.4 m s^−1^, and Δ*t*=0.55 [0.03] s at *U*^W^=0 m s^−1^). Thus, sidewinds negatively affect the landing time directly after take-off, whereas bumblebees landing from free-flight can fully compensate for the detrimental effects of sidewinds on travel time.

## Discussion

Winds are an ubiquitous characteristic of the natural environment of foraging bumblebees (Crall et al., 2017). Here, we studied how bumblebees perform landing maneuvers in steady sidewinds by recording 19,421 landing approaches of bumblebees towards a vertical surface in six different sidewind levels (0 to 3.4 m s^−1^), corresponding to conditions in nature (Crall et al., 2017; Riley et al., 1999).

### Landing strategy of bumblebees in steady sidewinds

We found that, in all tested wind conditions, bumblebees kept the optical expansion rate *r* approximately constant for brief periods of time during landing (Figure 5). We call such periods constant-*r* phases, and the corresponding constant-*r* value as the expansion rate set-point *r*\*. Bumblebees tended to stepwise increase *r*\* during the landing approach. This trend of increasing *r*\* with reducing distance is captured by a linear relationship with average slope *m*=−0.843 [0.01] between their logarithmic transformations, independent of sidewind speed (Figure S2; Table S4). Rate *m* resembles to the previously observed value for bumblebees landing in quiescent air on different landing platforms and at light intensities ranging from dusk to overcast daylight (Goyal et al., 2021), suggesting that the underlying control mechanism is conserved for a wide range of environmental conditions.

Rate *m* is equivalent to the time-to-contact-rate 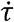 for the landing of birds (Goyal et al., 2021; Lee et al., 1991), and its magnitude in bumblebees is strikingly similar to that of landing pigeons 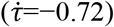 (Lee et al., 1993), hummingbirds 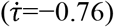 (Lee et al., 1991) and mallards 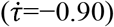 (Whitehead, 2020). Despite the similar slopes for birds and bumblebees, they differ substantially in landing strategy. Birds continuously increase *r* with reducing distance to the surface, whereas bumblebees do it in a stepwise manner.

The stepwise modulation of *r*\* causes landing bumblebees to fly at a range of optical expansion rates while converging towards the new set-point – a typical attribute of a step-response (Ogata, 2010). In still air, these time-evolutions, captured as entry segments, are the transient response of a sensorimotor control system that regulates the optical expansion rate (Goyal et al., 2022). The accelerations (or decelerations) during these entry segments bring the optical expansion rate closer to the desired set-point. Here, we showed that in a sidewind, bumblebees exhibit similar transient flight behaviors when switching between different optic expansion set-points. In addition to the stepwise variation of *r*, landing bumblebees also regularly generated low velocity flight phases, in which they hover or even briefly fly away from the surface.

### How sidewinds affect the landing dynamics and performance of flying bumblebees

#### (1) Sidewinds augment the transient flight responses of landing bumblebees

Here, we showed that bumblebees landing in sidewinds exhibited faster transient responses, expressed as optical expansion-acceleration 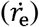 and faster mean body accelerations towards the landing platform (*Ā*_e_). This holds for all covariates values that influence the transient response of bumblebees, including distance from the landing surface (*y*_0_), required stepchange in optical expansion rate (Δ*r*) and the associated *r*\*. The higher 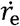 and *Ā*_e_ values during the transient response indicate that bumblebees generate higher control forces. Hence, the airspeed measuring mechanosensory modality of bumblebees affects aerodynamic force control during entry segments.

To assess this effect, we tested how *Ā*_e_ varied with the mean airspeed 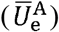 that bumblebees experienced during entry segments in different wind conditions. A linear fit captured this interaction well (slope = 0.204 [0.015] s^−1^, coefficient of determination *R^2^*= 0.98; Figure 9A). Thus, mean accelerations during entry segments increase approximately linearly with the airspeed induced by the sidewind. This linear relationship indicates positive feedback with a constant gain from the airspeed measuring mechanosensory modality to the vision-based flight regulator of landing bumblebees (Figure 9B). A similar multi-sensor feedback architecture was suggested for free-flying *Drosophila* (Fuller et al., 2014).

**Figure 9.**
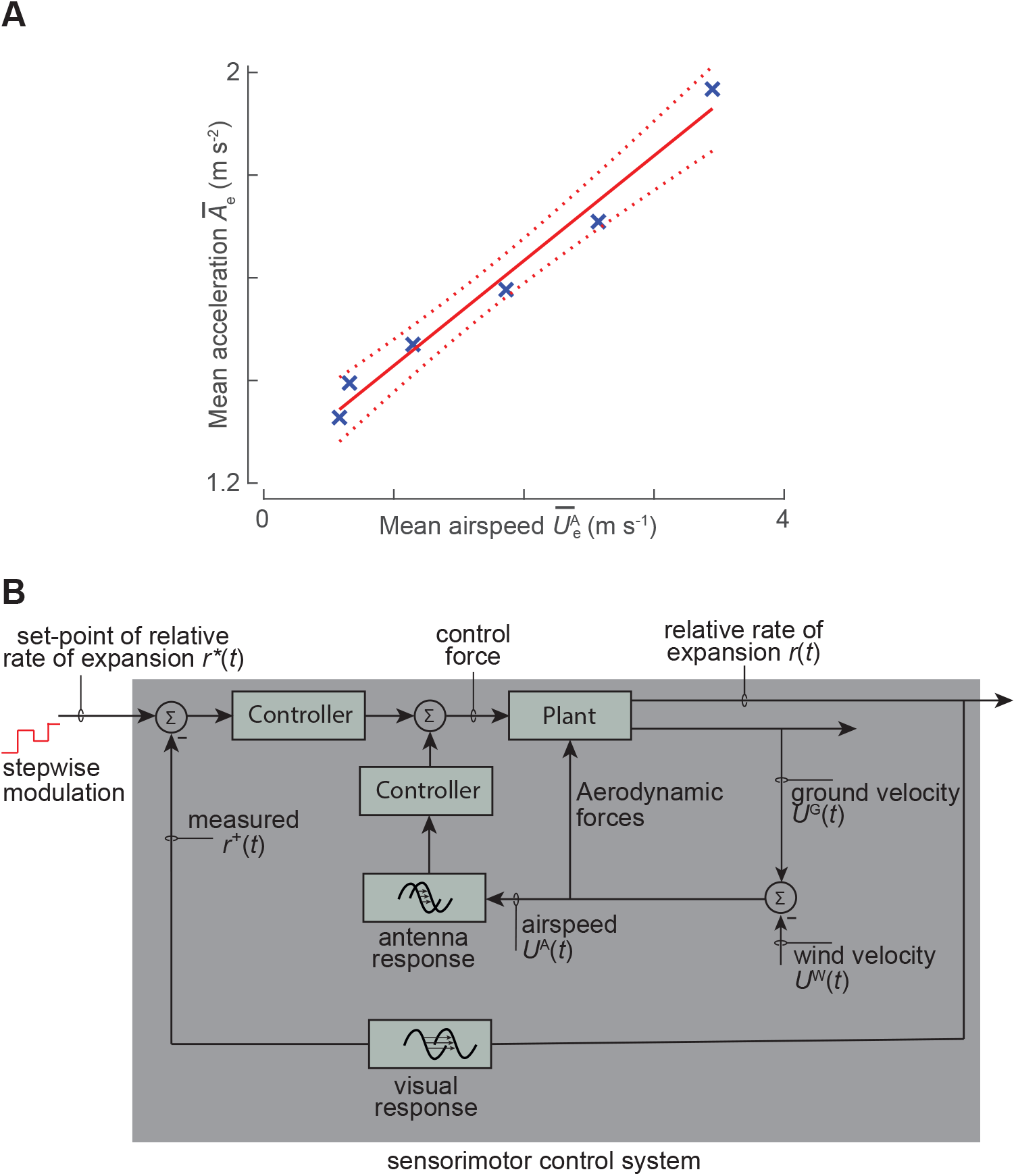
Proposed multimodal sensorimotor control system for bumblebees landing in sidewind, with airspeed measurement integrated in the visual-feedback loop. (A) During entry segments, the mean acceleration *Ā*_e_ increases approximately linearly with mean airspeed 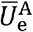, as estimated using our linear mixed model. Blue crosses depict the model results at each tested windspeed, and the solid and dotted red lines shows the linear fit and 95% confidence interval, respectively (see Materials and Methods). (B) Proposed model of multimodal sensory integration in landing bumblebees that explains the linear increase of the entry segment approach acceleration *Ā*_e_ with airspeed 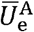. Bumblebees might use their antennae to measure the wind-induced mechanosensory input (airspeed) and integrate it with positive feedback in their vision-based sensorimotor control loop (Figure 1C). This fast positive feedback can provide active damping that counteracts the unstable oscillations of a visual feedback loop.

Mechanoreceptors on the bumblebee’s antennae (Jakobi et al., 2018; Taylor and Krapp, 2007) likely detect the changes in airspeed. As the neural processing time of information from antennae (~20 ms) is much shorter than that of the visual system (~50–100 ms), positive feedback from the antennal system can provide active damping to any vision-based regulator (Fuller et al., 2014). During sudden disturbances such as wind, active damping can reduce the oscillations of a slow visual feedback loop, as observed in free-flying *Drosophila* (Fuller et al., 2014). Active damping stabilizes the dynamics of insect locomotion in multiple other scenarios (Cheng and Deng, 2011; Cowan et al., 2006; Dyhr et al., 2013; Elzinga et al., 2012; Hedrick, 2011; Hedrick et al., 2009; Sun, 2014; Taylor et al., 2013).

Based on these observations, we propose that the airspeed sensing mechanosensory modality provides positive feedback – with constant gain – to the vision-based control of landing bumblebees (Figure 9B). During sudden wind disturbances, this fast mechanosensory feedback would reduce the oscillations caused by a slow visual control loop.

#### (2) Sidewinds increase the hover frequency in landing maneuvers

More hover phases could result in a longer landing duration, which in turn can negatively impact their foraging efficiency (Balfour et al., 2021). Why do bumblebees nevertheless tend to increase the hover phase number in faster winds? The increase in the transient response of the sensorimotor control system with wind velocity, independent of all other covariates, is analogous to bumblebees operating their visual feedback loop at a higher gain in still air. This increased gain in the *r*-based control loop can result in instabilities occurring at distances further away from the landing surface (Croon and De Croon, 2016). Thus, the characteristic hover phases exhibited relatively close to the landing surface in still air will occur on average at larger distances in the presence of winds. This is consistent with our findings that (a) independent of wind speed, hover frequency increased with decreasing distance from the landing surface, and (b) hover frequency increased with increasing wind velocity, independent of distance. This suggests that the hover phases in landing bumblebees are caused by the interaction of airspeed feedback with the vision-based control loop.

#### (3) Sidewinds increase the optic expansion rate set-points of landing bumblebees

Our analysis showed that landing bumblebees exhibit higher values of *r*\* in faster winds (Figure 5F), and thus the approach flight speeds during the constant-*r* segments are increased in high sidewinds. This resembles the dynamics found for honeybees landing in a headwind, where the mean set-point of translational optic flow increases with headwind speed (Baird et al., 2021), suggesting that the airspeed measuring mechanosensory modality, in addition to the sensorimotor control system, also influences the optic expansion rate set-points that landing bumblebees converge to.

#### (4) Sidewind increases the travel time of landing maneuvers, but only of landing after take-off

We used travel time of landing approaches as a metric for the landing performance of foraging bumblebees. For landings from free-flight, sidewinds did not affect these times, whereas for landings initiated directly after take-off travel times increased with sidewind velocity (Figure 8A and 8B, respectively).

Landings performed directly after take-off represent the rapid consecutive landings made by bumblebees when visiting flowers in a single flower patch. Our results contrast with those obtained for honeybees, where an increase in wind speeds did not affect the inter-flower flight duration (Hennessy et al., 2020; Hennessy et al., 2021). However, in these studies flowers were situated closer to each other than the platform distance in our study, and that the effect of headwinds in the wakes of the flowers was presumably lower and more variable than the uniform sidewinds in our study.

A second metric for landing performance is the average approach velocity 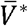 estimated using the average-track-based analysis method (Figure 4), which shows strikingly similar results as the travel time analysis. The observed 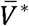 of bumblebees landing from free flight was constant for all tested wind conditions, whereas for landings directly after take-off 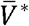 decreased with increasing sidewind speed (Figure 4G and 4H, respectively). Hence, the average-track-based analysis method is useful to estimate landing performance metrics such as the average approach velocity. However, it does not capture the observed complex kinematics of bumblebee landing maneuvers.

#### (5) How bumblebees compensate for the detrimental effects of landing in a sidewind

Both landing performance metrics travel time Δ*t* and 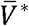 are thus the result of a complex interaction between decelerations towards the landing platform during constant-*r* phases, accelerations during entry segments, and hover phases.

The occurrence of hover phases increases with sidewind speeds, for all tested conditions (Figure 7). Hovering increases Δ*t*, and thus it can directly explain the increase in Δ*t* with wind speed, for landings directly after take-off (Figure 8B). In contrast, Δ*t* of landing approaches from free-flight do not differ between wind conditions (Figure 8A), whereas their hover frequency does increase with wind speeds (Figure 7B).

How do bumblebees landing from free flight in high winds compensate for the detrimental effect of the concomitant increase in hovering? Bumblebees landing in sidewind integrate airspeed measurements in the vision-based landing control loop (Figure 9). By doing so, landing bumblebees increased their average optic expansion set-point with increasing wind speed (Figure 4G), and they performed more rapid transient flight responses at higher windspeeds leading to faster approach accelerations (Figure 6E,F). This landing control mechanism was similar between bumblebees landing from take-off or from free-flight, as all related control parameters (*r*\*, *V*\*, 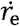, *Ā*_e_) did not differ between these landing types. However, the hover probabilities differed between landings from take-off and free-flight: the effect of wind on hovering probability was on average 88% larger for landings from take-off. This lower hover frequency in landings from free-flight explains thus why for these landings bumblebees were able to fully compensate for the negative effects of increased hover frequencies at high winds, but bumblebees landing after take-off were not.

### Energetic costs of landing in a sidewind

In our set-up, landings directly after a take-off took more time at higher wind speeds. This can negatively influence foraging efficiency when visiting multiple flowers within a flower patch (Balfour et al., 2021). Moreover, bumblebees foraging in the fastest tested windspeed landed 70% less than in still air. Similar dynamics were observed in field and semi-field conditions, where the winds negatively impacted the foraging rate of honeybees (Hennessy et al., 2020; Hennessy et al., 2021; Pinzauti, 1986; Vicens and Bosch, 2000).

These combined results suggest that wind can negatively affect the fitness of individual insect pollinators and their colony (Riessberger and Crailsheim, 1997), and the efficacy of their pollination services (Tuell and Isaacs, 2010). Understanding these effects is crucial as insect pollinators support biodiversity (Ollerton et al., 2011) and global food production (Klein et al., 2007). This is even more pertinent with a predicted increase in windspeeds due to climate change in several areas of the world (Hosking et al., 2018). Future work in this direction could address the direct effect of winds on the colony fitness and pollination dynamics.

### Conclusion

Wind is an important yet understudied factor that influences the control strategy of flying insects. Here, we show how landing bumblebees use visual and airflow cues to advance towards a landing surface in steady lateral winds, and how they compensate for detrimental wind effects on their landing performance. We showed that bumblebees in these winds approach a landing surface using a visually-guided strategy similar to that in still air, but with some key differences. Like landings in still air, in a sidewind bumblebees advanced towards the landing surface by regulating optical expansion rate and exhibiting bouts of acceleration and deceleration phases, along with occasional hover phases. Bumblebees landing in a sidewind travelled faster towards the landing surface than in still air, but exhibited more hover phases. The occurrence of more hover phases can reduce foraging efficiency as these tend to increase the landing time. But, by travelling faster towards the surface, bumblebees fully or partly compensated for this hover-induced increase in travel time, depending upon whether they landed from a free-flight or after a take-off, respectively.

Based on our observations, we suggest that landing bumblebees implement a positive feedback from their airspeed measuring mechanosensors to their visual feedback loop, as previously proposed for flying *Drosophila* (Fuller et al., 2014), which provides active damping to unstable oscillations of the visual feedback loop in the event of external disturbances.

In nature, winds directly and indirectly influence the landing dynamics of bumblebees. The direct influence corresponds to the effects of mean wind speeds and the fluctuations around these speeds on the landing dynamics. The indirect influence corresponds to the impact on the visual information that bumblebees perceive due to swaying of flowers in winds (Hennessy et al., 2020; Kapustjansky et al., 2010). Our study of the effects of mean winds on the landing dynamics of bumblebees is a step towards better understanding the exemplary ability of bumblebees to mitigate the detrimental effect of wind on flight control.

## Acknowledgements

We thank Antoine Cribellier, Cees Voesenek and Wouter van Veen for useful discussions, Remco Pieters for his help in building the experimental set-up, and Remco Huvermann for providing the bumblebees.

## Data and code availability

The data and code used for this study are publicly available at https://data.mendeley.com/datasets/mww9m8r3dk/draft?a=f98651c6-e4e3-4f1b-994d-4edd10ec6552 and https://github.com/kaku289/nimble-bbee-analysis/tree/landing_steady_winds, respectively.

